# Sequential replay of non-spatial task states in the human hippocampus

**DOI:** 10.1101/315978

**Authors:** Nicolas W. Schuck, Yael Niv

**Author notes:** Corresponding author contact: Max Planck Research Group NeuroCode Max Planck Institute for Human Development Lentzeallee 94, 14195 Berlin, Germany.

## Abstract

Neurophysiological research has found that previously experienced sequences of spatial events are reactivated in the hippocampus of rodents during wakeful rest. This phenomenon has become a cornerstone of modern theories of memory and decision making. Yet, whether hippocampal sequence reactivation at rest is of general importance also for humans and non-spatial tasks has remained unclear. Here, we investigated sequences of fMRI BOLD activation patterns in humans during wakeful rest following a sequential non-spatial decision-making task. We found that pattern reactivations within the human hippocampus reflected the order of previous task state sequences, and that the extent of this offline reactivation was related to the on-task representation of task states in the orbitofrontal cortex. Permutation analyses and fMRI signal simulations confirmed that these results reflected underlying BOLD activity, and showed that our novel statistical analyses are, in principle, sensitive to sequential neural events occurring as fast as one hundred milliseconds apart. Our results support the importance of sequential reactivation in the human hippocampus for decision making, and establish the feasibility of investigating such rapid signals with fMRI, despite its substantial temporal limitations.

**Highlights:**

- We provide fMRI evidence for sequential pattern reactivation in the human hippocampus
- Sequences of patterns reflect states from a sequential, non-spatial decision-making task
- Simulations show that our novel fMRI analysis is sensitive to fast sequences of sub-second neural events
- Results support the importance of sequential reactivation in the human hippocampus for decision making

## Introduction

The hippocampus plays an important role in memory^1–3^, and is known to represent spatial as well as non-spatial information that is relevant to an animal’s current location within a ‘map’ of the ongoing task^4–8^. Recent research has suggested that hippocampal memories are also used to inform spatial decision making and planing by reactivating neurally encoded experiences that are relevant for the current task^9,10^. Specifically, studies in rodents have shown that hippocampal representations of spatial locations are reactivated sequentially during short on-task pauses, longer rest periods, and sleep^11–13^. This sequential reactivation, or replay, is related to better planning^12^ and memory consolidation^14^, and suppression of replay-related short-wave ripples impairs spatial memory^15^.

While these findings have provided insights into the hippocampal computations underlying spatial decision making in animals, what role replay plays in non-spatial decision making tasks in humans has remained unclear. We instructed participants to perform a sequential non-spatial decision making task, and recorded functional magnetic resonance imaging (fMRI) activity during resting periods before and after the task. The sequential nature of the task was critical to correct performance, ensuring that participants would encode sequential information while completing the task. We investigated whether sequences of fMRI activation patterns during rest reflected hippocampal replay of task states. Evidencing such replay, transitions between hippocampal fMRI activity patterns were related to sequences of task states. Careful analyses and simulations confirmed that the structure seen in hippocampal data reflects sequential information in the BOLD responses above and beyond any structure introduced by state classifiers trained on task states. Importantly, reactivation in the hippocampus during rest was associated with better representation of the same task states in the orbitofrontal cortex during decision making, which in turn was related to better performance of the task, in line with our previous work^16^.

Our results demonstrate sequential reactivation of non-spatial decision-making states in the human hippocampus and suggest that the interaction between hippocampal and prefrontal brain systems supports the construction and use of representations reflecting the structure of the current task. Our findings, together with a set of rigorous statistical tests and simulations, also establish the utility of noninvasive fMRI to detect possibly rapid replay events, despite the low temporal resolution of this method.

## Results

Thirty three human participants performed a sequential decision-making task that required integration of information from past trials into a mental representation of the current task state^16^ (see Methods). Specifically, each stimulus consisted of overlapping images of a face and a house and participants’ main task was to make age judgments (old or young) about one of the images (Fig. 1A). The category to be judged (face or house) was instructed before the first trial. Subsequent category switches were determined by the following rule: if the age in the current trial was the same as the age in the previous trial, then the judged category remained the same; on the other hand, if the age on the current trial was different from the age on the previous trial, the participant had to switch to the other category from the next trial onward (Fig. 1B). This created a ‘miniblock’ structure where each miniblock involved judgment of one category. No age comparison was required on the first trial after a switch. Miniblocks were therefore at least two trials long, and on average lasted for three trials. These task rules resulted in a total of 16 ‘task states’ reflecting the ‘location’ in the current miniblock and were experienced in a structured order (Fig. 1C). For example, the (Ho)Fy state, indicating a young face trial that followed an old house trial, was only experienced at the end of a house (old) miniblock, as the first trial of the next face (young) miniblock. This particular structure meant that although the task was not spatial, it involved navigating through a sequence of states that had predictable relationship to each other, as in a virtual maze. Participants performed the task with high accuracy (average error rate: 3.1%, time-outs: 0.6%, reaction time: 969 ms), improving their performance throughout the course of the experiment (see Fig. 1D, significant linear trends of task block for reaction times and errors, both *p*’s<.001, see also Supplemental Information, SI, Fig. S4, for further details).

**Figure 1:**
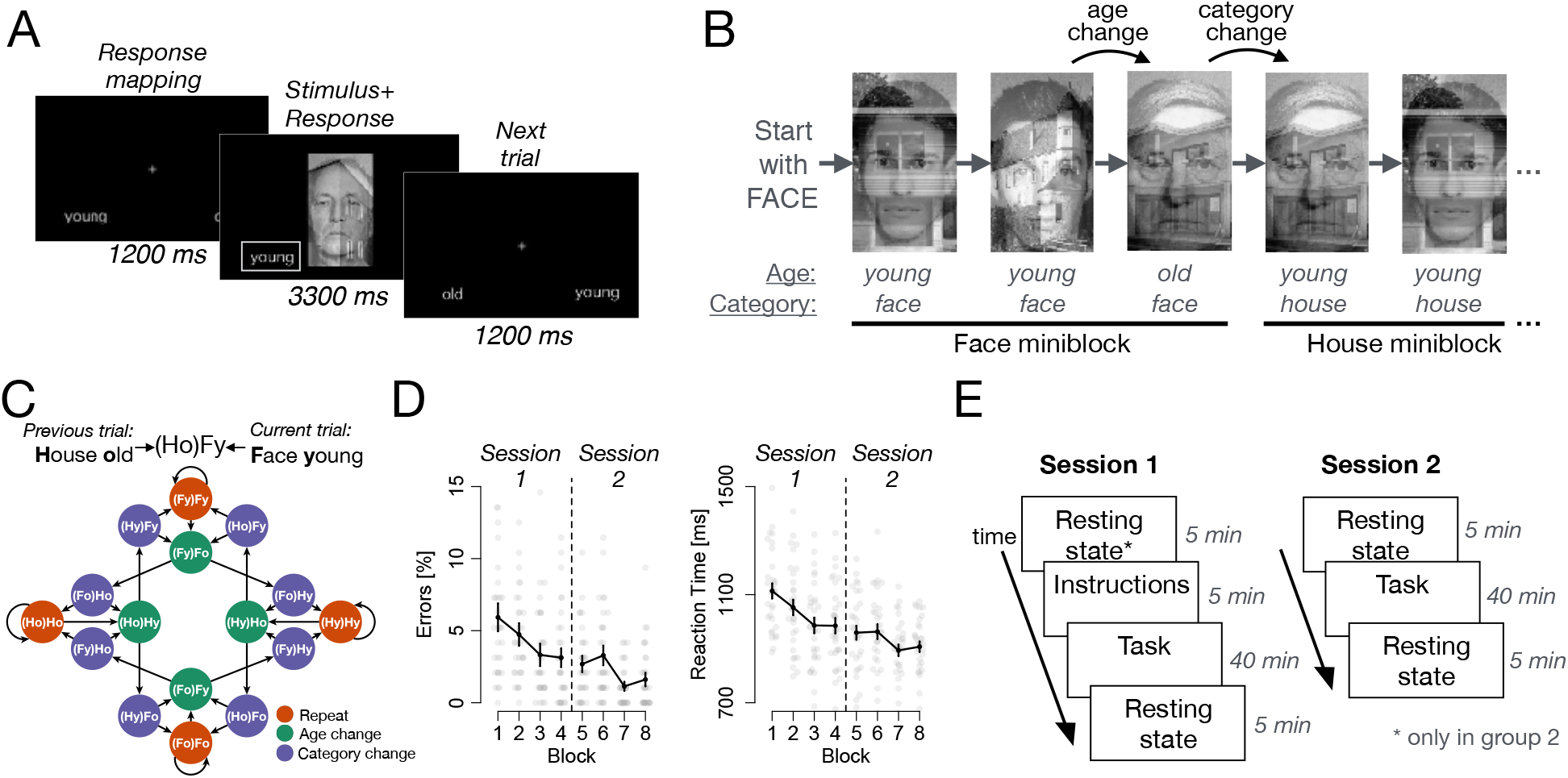
Experimental task and performance. *(A)*: On each trial, participants had to judge the age of either a face or a house shown overlaid as a compound stimulus. Trials began with the display of a fixation cross and the response mapping (how left/right was assigned to old/young; 1200ms), followed by the stimulus. Responses could be made at any time, and the stimulus stayed on screen for an average of 3300 ms. *(B)*: The task required participants to switch between judging faces and houses following each time the age changed between two trials, see text. *(C)*: The state space of the task, reflecting the abstract space which participants traversed, analogous to a spatial maze although non-spatial from the point of view of the participant. Each node represents one possible task state, and each arrow a possible transition. All transitions out of a state are equally probable, occurring with *p* = 0.5. Each state of the task is determined by the age and category of the previous and current trial, indicated by the acronyms (see legend). States are colored based on their ‘location’ within a miniblock: trials within a miniblock in which the age and category were repeated (orange), trials at the end of a miniblock in which the age changed (green), and trials entering a new miniblock where the category changed (purple). *(D)*: Average error rates and reaction times across the two sessions. Bars: ± 1 S.E.M. Grey dots represent individual subjects. *(E)*: The experiment extended over two sessions, each of which included about 40 minutes task experience flanked by resting state scans. *: The pre-task resting state scan in Session 1 was performed only for a subgroup of our sample (*N* = 10; group 2).

The experiment was comprised of two sessions during which participants engaged in the above decision-making task while undergoing fMRI. The first session included ~ 5 minutes of task instructions and four runs of task performance (388 trials, about 40 minutes duration). The second session took place one to four days later and was identical to session 1, but without instructions (Fig 1E). Resting state scans consisting of 5 minute periods of wakeful rest without any explicit task or visual stimulation (100 volumes per resting state scan) were administered for *N* =23 participants (group 1) after session 1, before session 2 and after session 2, resulting in a total of 300 whole brain volumes acquired during rest. A second group of participants (*N* = 10; group 2) underwent the same procedures as group 1, plus one additional resting state scan at the beginning of session 1, before having had any task experience (including being exposed to task instructions). This resulted in a total of 400 whole brain volumes acquired during rest. The analyses reported below focus on fMRI data recorded during these resting scans. Resting-state data acquired after participants had task experience will from hereon be referred to as the TASK rest condition, whereas resting state data acquired before the task as the PRE rest condition. Data recorded while participants received instructions will serve as another control and be referred to as the INSTR condition. To account for differences in the number of data points constituting the TASK vs control conditions, we used a size-matched TASK condition where appropriate. Notably, while none of these conditions involves active experience of the sequential decision-making task, they differ in whether the task has been experienced before or not, and therefore in whether hippocampal replay might be expected or not.

The main goal of our study was to investigate sequential reactivation, or replay, of task-related experiences in the human hippocampus during rest. To this end, we trained a multivariate pattern recognition algorithm (see Methods) to distinguish between the activation patterns associated with each of the 16 task states in the data recorded during task performance (Fig. 2A,B). Leave-one-run-out cross-validated classification accuracy on the task data from the hippocampus (HPC) was significantly higher than chance and than classification obtained in a permutation test (11.6% vs 7.1% in the permutation test, *t*_32_ = 8.9, *p* < .001, chance level is 6.25%, Fig. 2C), indicating that hippocampal activation patterns indeed reflected task states. We then applied the trained classifier to each volume of fMRI data acquired during the resting state scans. Although classification accuracy cannot be assessed for the resting scan data (due to lack of ground truth), we could assess the quality of the classification using the mean unsigned distance to the decision hyperplane, a proxy for classification certainty^17^. This distance was larger in the TASK condition compared to simulated spatiotemporally-matched noise (‘NOISE’, *t*_32_ = 12.9, *p* < .001; for simulation details see Methods and SI) and compared to the PRE condition (*t*_9_ = 2.1, *p* = .031, group 2 only, Fig. 2D). This suggests that fMRI patterns recorded during resting-state scans following task experience could reflect reactivation of task states, in line with previous findings^18–20^.

**Figure 2:**
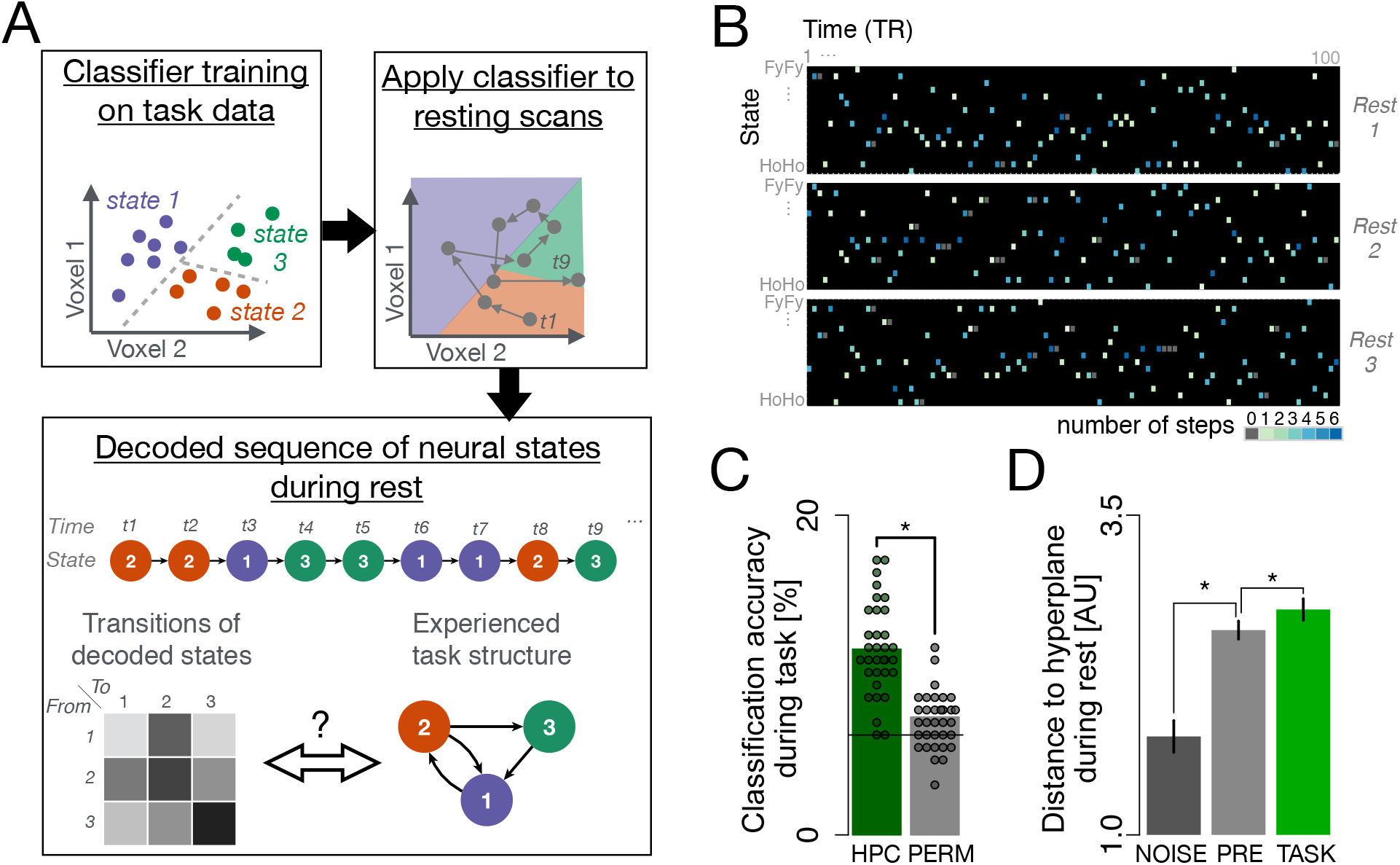
Sequential replay decoding analysis. *(A)*: Illustration of analysis procedure. For simplicity, only two dimensions and three state classes are shown. We first trained a classifier to distinguish between the different task states in the hippocampal fMRI data acquired during the task. The trained classifier was then applied to each volume of fMRI data recorded during resting sessions (grey dots). This resulted in a sequence of predicted classifier labels that was then transformed into a ‘transition matrix’ *T* that summarized the frequency of decoding each pair of task states consecutively. The structure of the decoded sequences, as summarized by this matrix, was then compared to the sequential structure of the task (see text). Note that the real analysis involved 16-way classification of >1000 dimensional data, which was compared to the task state space shown in Fig. 1C. *(B)*: Example data from one randomly selected participant. Each dark rectangle illustrates the sequence of classified states for the 100 volumes of fMRI data recorded in one resting state scan (depicted are three resting state scans acquired throughout the experiment; see Fig. 1E). Columns represent time, and rows states. Each color-filled cell represents the state classified at the respective time point, and color indicates the distance (in steps in the state space; Fig. 1C) from the state decoded in the previous timepoint (i.e., the previous TR). *(C)* Classification accuracy during task performance was significantly higher in hippocampal data (HPC) than in a permutation test (PERM). The solid line represents the theoretical chance baseline of 100/16=6.25. *(D)*: Average distance to the hyperplane for classified states during rest in the NOISE (dark grey, left bar), PRE (light grey, middle bar, N=10) and TASK conditions (green, rightmost bar, N=33). Larger distance indicates higher certainty in the classification of the state. Each dot indicates one participant, bars within-subject S.E.M., *: *p* < .05.

The defining aspect of replay is that previously experienced states are reactivated *sequentially*. We therefore asked next whether it is theoretically possible to measure rapid sequential replay events (on the order of few hundreds of milliseconds in humans^21^) using fMRI, given its low temporal resolution. To this end, we simulated fMRI activity that would result from fast replay events (see SI and below), and asked what order and state information could be extracted from these spatially and temporally overlapping patterns. The slow hemodynamic response measured in fMRI causes brief neural events to impact the BOLD signal over several seconds. Although these same dynamics might obscure the details of a replayed sequence, our simulations showed that two successive fMRI measurements could still reflect two states from the same sequence, for instance the first and last element of a multi-step replay event (see SI). Because replay events mainly reflect short sequences of states^13^ (figure 3C in Ref. 13), if the activity we measured in the hippocampus at rest indeed reflects sequential replay, we can therefore expect that consecutively decoded states be close in the task’s state space (that is, separated by few intervening states in Fig. 1C). This is analogous to the expectation that spatial replay events would lead to sequential activation of place cells with place fields nearby in space – even if some place cells were erroneously not identified as being active. However, given the low accuracy of correctly decoding task states during task performance, could we even expect to successfully decode a pair of states from the same replay event? Our simulations showed that we could: because brain activity recorded after a rapid replay event presumably includes several superimposed states (Fig. S5B), the likelihood of classifying *one out of several* replayed states in each resting state brain volume is actually considerably higher than the overall decoding accuracy when classifying a single prolonged event during task. Our theoretical analysis showed that the chance that analyzing two consecutive brain volumes results in decoding one (ordered) set of two states out of several possible sets caused by the same replay event may be on the order of the overall decoding accuracy (~10%; see SI).

**Figure 3:**
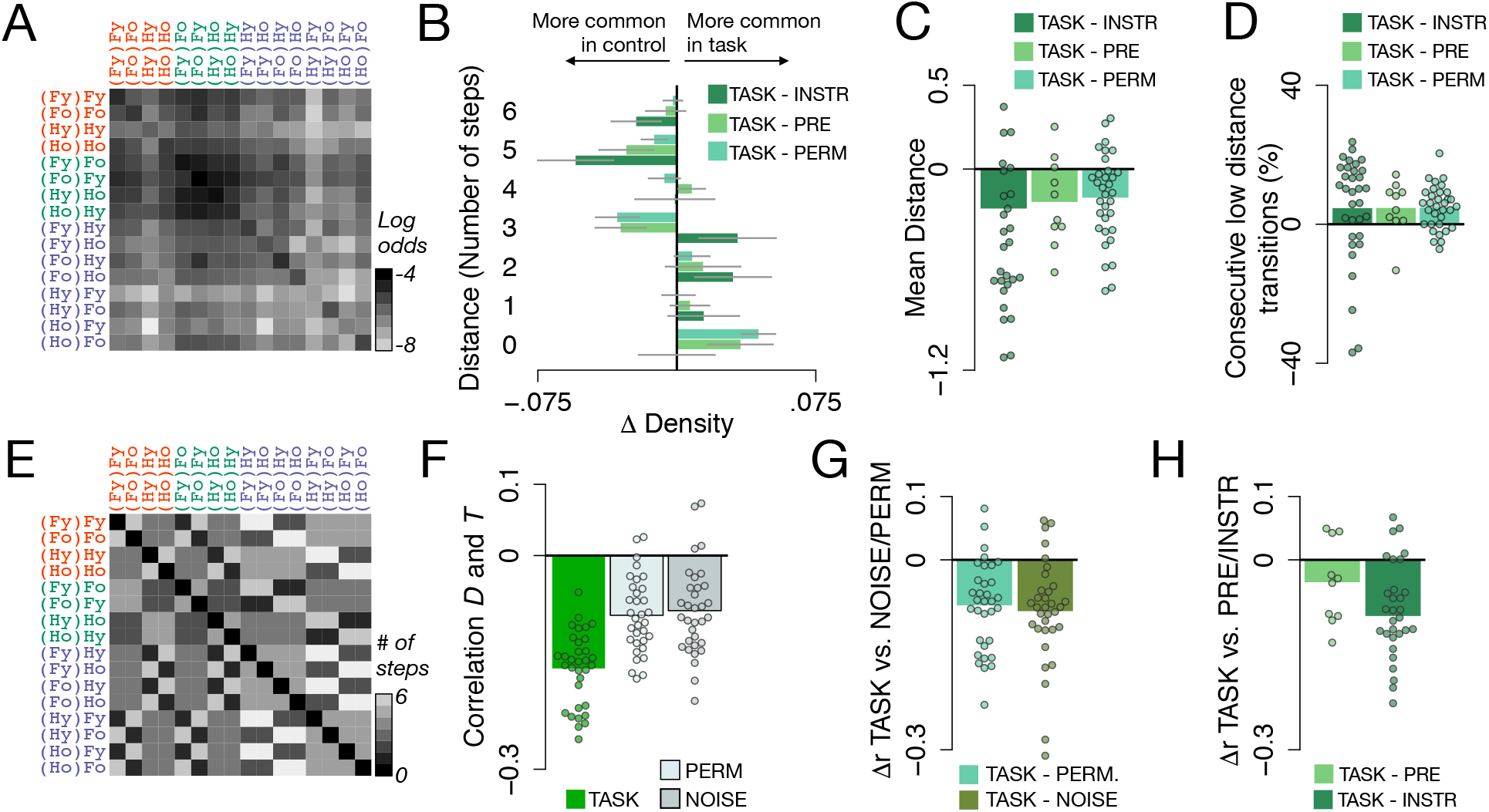
Hippocampal state transitions during rest are related to state distances in the task. *(A)*: The matrix *T*, expressing the log odds of transitions between all states in the sequence of classification labels in the hippocampal TASK rest data, averaged across all participants. Y-axis: first state, x-axis: second state, in each consecutively decoded pair. Darker colors reflect a higher probability of observing a pair in the data. *(B)*: Relative distributions of number of steps separating two consecutively decoded states. A distance of 1 corresponds to a decoded state transition as experienced in the task, 2 corresponds to a transition with one item missing in between as compared to the task, etc. Barplots show the difference in relative frequency (Δ Density) with which each transition type was observed in the TASK condition compared to INSTR and PRE control conditions and a permutation test (PERM), see legend. Smaller distances are more frequently observed in the TASK data, whereas larger distances are more common in the control data, suggesting that the TASK resting-state data reflect reactivation of short sequences *(C)*: The average distance in state space of two consecutively decoded states was significantly lower in the TASK data as compared to the INSTR, PRE and PERM controls (all *p*s <.05, t-test comparing difference to 0). *(D)*: Low-distance transitions (fewer than 3 steps) occurred in succession significantly more frequently in the TASK data compared to all controls (all *p*s <.05). *(E)*: The matrix *D*, indicating the minimum number of steps between each pair of states in the task (i.e. the state distances). Lighter colors reflect larger distance between states. *(F)*: Average correlations between the state distance matrix *D* and the corresponding decoded transition matrix *T* in the TASK condition (green bar, left), as compared to a permutation test (PERM; light grey, middle) or when the same classifier was applied to participant-specific spatio-temporally matched noise (NOISE; dark gray bar, right, see also Fig. S1). *(G)*: Within-participant differences between correlations in TASK versus the PERM and NOISE controls (all *p*s <.05) *(H)*: The anti-correlation between *D* and *T* in the PRE and INSTR phases was lower than in the TASK resting state sessions (matched in amount of data compared). Dots reflect differences in correlations for individual participants.

Having established that, in principle, we can detect sequential replay in fMRI data, we next investigated whether the sequences of states we decoded in the TASK resting data (Fig. 3A) reflected the sequential structure of the experienced task. We note up front that because the classifier was trained on task data that were themselves sequential, signs of sequentiality of classifier predictions might arise even in random noise—although clearly those data do not reflect sequential replay. We therefore conducted a series of carefully controlled assessments of the levels of sequentiality in our data to ensure that we were detecting true sequential replay, and not merely unveiling the properties of the classifier. Indeed, we found in our data several forms of sequentiality that are predicted by replay, above and beyond what we could find in a series of closely matched controls.

First, we predicted that replay would be reflected in a smaller number of steps that separate two consecutively decoded states, as indicated by the above-mentioned simulations. In line with this idea, the number of steps between state transitions decoded in the TASK resting condition was smaller, on average, than the distance between states in the INSTR condition (*t*_32_ = 2.4, *p* = .01), smaller than the distance found in the PRE condition (*t*_9_ = 2.3, *p* = 0.02, group 2 only) and smaller compared to a permutation test in which classified states were randomly reordered to control for overall state frequency (PERM condition: *t*_32_ = 4.6, *p* < .001; Fig. 3B,C). Second, because replay events are separated by long pauses^21^, and sequentiality should be present only following the replay events, we expected the occurrence of short-distance state pairs to be clustered in time. Indeed, short-distance state pairs (less than 3 steps apart) were not only more frequent than expected, but were also more likely to occur in clusters in the TASK rest condition compared to the INSTR (*t*_32_ = 1.7, *p* = .046), PRE (*t*_9_ = 1.9, *p* = .044, group 2 only), and PERM controls (*t*_32_ = 4.5, *p* < .001, Fig. 3D). Next, we ensured that the above results could not be explained by sustained state activation, or only one particular decoded state distance. To this end, we removed state repetitions (“self transitions”) from the data and tested whether the prevalence of each step size (a transition between two states separated by *k* steps) was linearly related to the distance between the two states in task space. In other words, we tested whether the empirical frequency of decoding each pair of task states consecutively (the “transition probability” for each pair of decoded states, summarized in matrix *T*; Fig. 3A) was negatively correlated with the distance *D* between the states in the task (where *D_ij_* corresponds to the minimum number of steps necessary to get from state *i* to state *j*; Fig. 3E). The correlation between *T* and *D* was indeed significantly negative (average *r* = −.16, *t*_32_ = −17.7, *p* < .001, t-test of individual correlations across participants, Fig. 3F), and was substantially more negative than the correlation seen in the PERM control (*r* = −.08, *p* < .001, reflecting an effect of overall state frequency; Δ*r* = −.07, *t*_32_ = −5.8, *p* < .001). Applying the trained classifier to matched fMRI noise (NOISE control, see Methods and SI, Fig S1) also resulted in a negative correlation (*r* = −.08, *p* < .001, showing that temporal contingencies between states in the classifier training data can lead to spurious correlations), which was nevertheless significantly weaker than the correlation found in the TASK rest data (Δ*r* = −.08, *t*_32_ = −5.6, *p* < .001, Fig. 3G). Importantly, our hypothesis that sequential reactivation of task-state representations during rest was caused by task experience was also supported by a significantly stronger anti-correlation between *T* and *D* in the TASK condition as compared to the INSTR data (*t*_32_ = −12.1, *p* < .001, when comparing a subset of the TASK condition matched in number of TRs to the INSTR data), as well as the PRE resting scan (*t*_9_ = −7.9, *p* < .001, group 2; *p* = .059 when compared to only the first resting scan in TASK condition), as shown in Fig. 3H. Finally, we also assessed the effect of the sequential structure in the training data on our results in an additional control analysis in which we systematically excluded sets of state pairs from classifier training (see SI, Fig. S2), to test if, as a result, these pairs would seem to have a lower transition frequency in the resting data. This concern was allayed as the excluded transitions were observed as often as the included transitions (*t*_32_ = 0.3, *p* = 0.73), in line with our conclusion that the transition frequencies observed during rest reflected sequential reactivation above and beyond any sequential structure in the classifier.

In order to investigate the effects of task experience on pair-decoding frequency data *T* while simultaneously (a) excluding state repetitions, (b) controlling for the above-mentioned effect of temporal contingencies in the classifier training and (c) incorporating the different sources of between- and within-participant variability, we performed a logistic mixed-effects analysis. In this, we modeled both the effect of interest (*D*) and nuisance covariates that could potentially affect *T* (see Methods). We will henceforth call the effect estimate (beta weight) of the distances *D* on the data *T* in this model ‘sequenceness,’ and the nuisance effects ‘randomness.’ Comparing a model of the frequency of transitions *T* that contained only randomness regressors with a model that also included the sequenceness (task distances) regressor *D*, we found no difference in model fit in the PRE rest condition (Aikaike Information Criterion, AIC, 3651.4 vs. 3651.5 for the model without and with the sequenceness regressor, respectively, 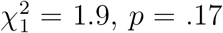). In the TASK rest condition, in contrast, adding the sequenceness regressor improved model fit significantly (AIC 3645.4 vs. 3642.1, lower numbers indicate a better fit, 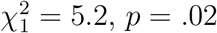, group 2 only and considering only the first TASK resting scan from the first session to equate power with the PRE analysis above). Indeed, the difference between the two datasets was statistically significant: including both PRE and TASK conditions within one model showed improved fit when the interaction of condition factor with sequenceness and randomness was included (AIC 3660.2 vs. 3674.1, *p* < .001). Figure 4A,B shows the sequenceness and randomness effects in the TASK compared to the PRE condition. Comparing the INSTR to the TASK condition in all participants showed the same pattern of effects: No effect of the sequenceness regressor was found in the INSTR condition (AIC 10046 vs 10047, *p* = .27), but there was a significant effect in the TASK rest condition (AIC 10130 vs. 10146, *p* < .001, TASK data matched in size to equate power), see Figure 4C,D. However, here the combined model indicated no interaction between condition and sequenceness vs. randomness (10142 vs 10130, *p* > .1). Note that the lack of sequenceness before task experience shows that our modelling analysis successfully controlled for bias effects due to the temporal contingencies between states in the classifier training data. Analyzing data from all participants (groups 1 & 2) and all TASK resting-state scans with this model indeed showed that the inclusion of a state distance factor led to significantly better model fits even after controlling for the randomness (bias) effect as above (AIC 10789 vs 10780, 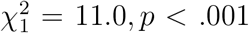), supporting the conclusion that previously experienced sequences of task states are replayed in the human hippocampus during rest periods. These results were unaffected by the choice of distance metric, see SI. No comparable pattern of results emerged when data from the orbitofrontal cortex, a brain area known to contain task-state information during decision making^16,22^, were analyzed in a similar fashion (see SI).

**Figure 4:**
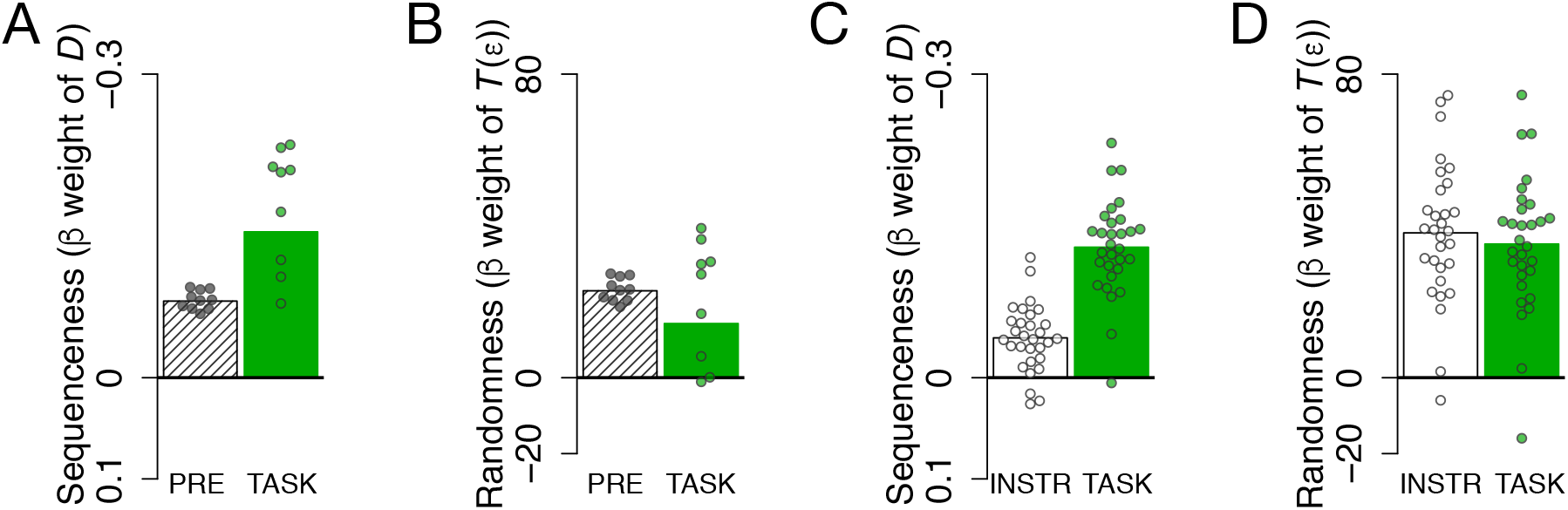
Effect of state distance (sequenceness) on transition frequency in hippocampal data is specific to TASK rest conditions. Bars indicate strength of fixed effects in mixed effects model (see text). Dots indicate individual random effects. Note that variability of dots in this case cannot be used to infer significant differences. *(A)*: Effect of sequenceness regressor *D* on resting data from the PRE and TASK conditions. Model comparisons based on AIC showed that the sequenceness regressor led to better model fit in the TASK but not the PRE condition. *(B)*: Effect of randomness across the PRE and TASK conditions. The randomness regressor *T*(*ϵ*) captures the sequentiality in the data due to classifier bias, see Methods. *(C)*: Sequenceness in the INSTR and TASK conditions, as in panel (A). Adding the sequenceness regressor led to better model fit only in the TASK condition. *(D)*: Randomness in the INSTR and TASK conditions as in panel (B).

The above analyses relied on the forward distance between states, as experienced during the task. We next tested whether the sequenceness found in the TASK condition could be explained better by alternative forms of replay, namely backward replay or forward replay of partial states such as the stimuli experienced. To this end, we defined alternative distance matrices corresponding to the above hypotheses, and tested the power of these alternative models to explain the sequences of states decoded during rest. For these analyses, instead of distances we used one-step task transition matrices to avoid statistical disadvantages of alternative models that have very evenly distributed distances (high entropy). As in our original analysis, all 1-step matrices were based on the task state diagram. The alternative 1-step matrices were created by either transposing the original 1-step matrix (backward analysis) or by assuming that only partial aspects of each trial’s state are represented, for instance by computing the experienced transitions between attended stimuli without representing the events in the previous trial (see Methods). As the classifier was trained to distinguish all 16 possible states, we assumed that different states corresponding for instance to the same stimulus would be fully aliased. We tested four alternative hypotheses by calculating the likelihood that the observed sequences of states were generated by (a) replay of states containing the stimulus on the current trial (‘stimulus model’, Fig. 5A), (b) replay of states containing only information about the currently attended category (‘category model’, Fig. 5B), (c) replay of states containing information about the attended category on the current and previous trial (‘category memory model’, Fig. 5C), and (d) backward replay of full states, in the opposite order as they were experienced (‘backward model’, Fig. 5D), and comparing these to the likelihood of the data being generated by forward transitions between full states (the one-step version of our original hypothesis; ‘full state model’, Fig. 5E). Model comparison using the same mixed effects models as above showed that the 1-step transitions assuming full state representation (Fig. 5E) led to a better model fit compared to all four alternative models (AIC: 20808, 20808, 20806, 20796, for the 4 alternative models, respectively; AIC of true model: 20782, see Fig. 5F).

**Figure 5:**
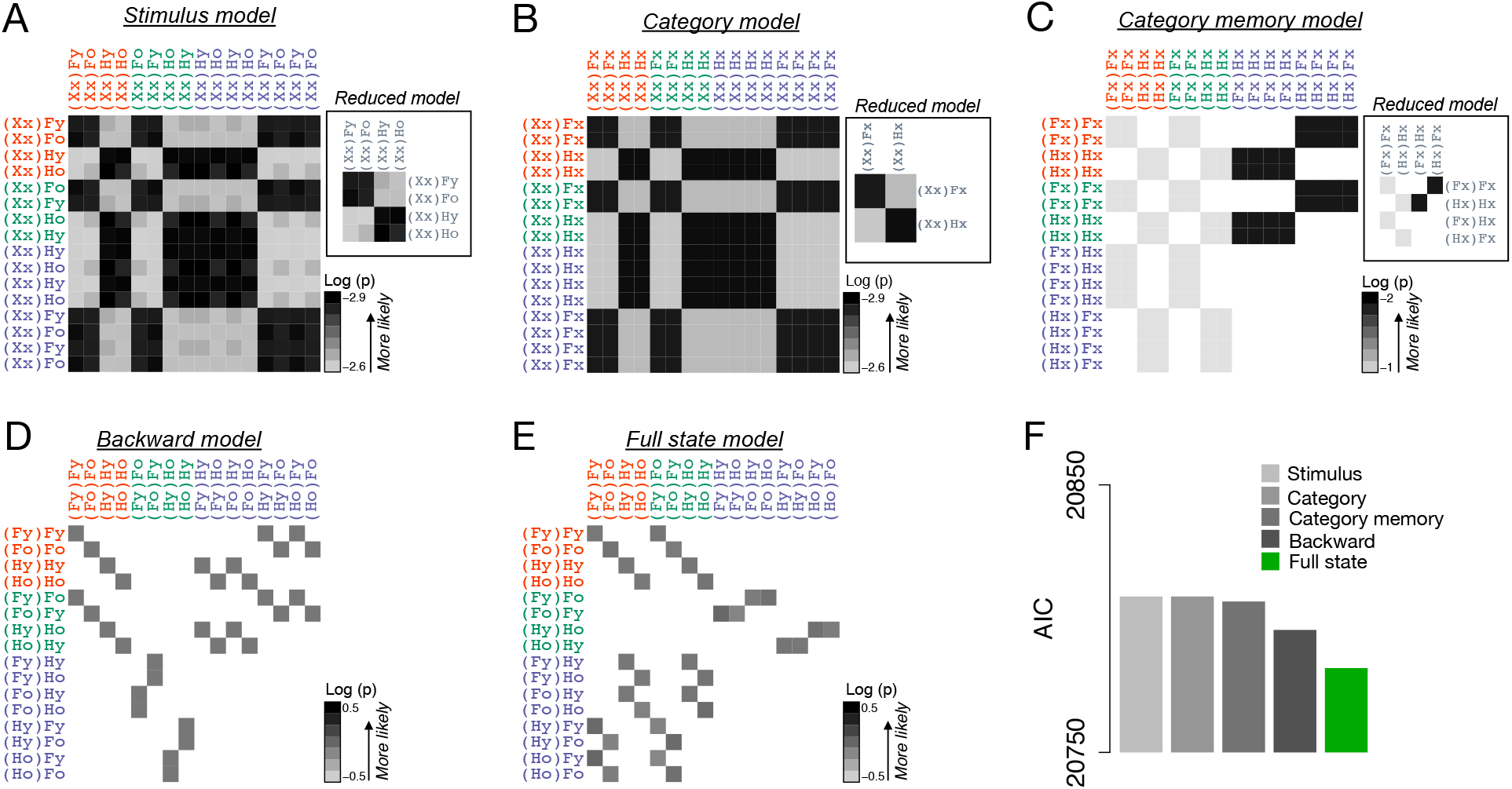
Alternative state transition matrices do not explain hippocampal state sequences during rest. *(A-E)* Alternative state transition matrices. Rows indicate origin states and columns indicate receiving states for a given transition, see text. Color shading indicates log likelihood of the corresponding 1-step transition under each alternative hypothesis, see legend and Methods. Empty (white) cells indicate that a transition is not possible. ‘Reduced model’ panels in A-C show the transition matrix when aliased states are collapsed. (E) Corresponds to the one-step transitions for our original hypothesis (compare to Fig. 3E). *(F)* Akaike information criterion (AIC) when data from the TASK rest condition were modelled using the transition matrices shown in A-E. The full state model explained the data best (lower AIC scores indicate a better model fit).

Similar to our above analyses investigating the theoretical possibility of decoding fast replay states from fMRI data, we also test whether the observed sequenceness in hippocampal fMRI data could have been caused by fast sequences of neural events in principle. For this we simulated fMRI signals generated by sequences of hypothetical neural events occurring at different speeds, and asked at which speed the above analyses can uncover the underlying sequential structure. In these simulations, each neural event triggered a hemodynamic response in a distributed pattern of voxels (see SI; Fig. S3). When the signal-to-noise ratio was adjusted to yield state-decoding levels that were matched to our data (12.1% accuracy in simulations, vs. 11.6% in the data), significant correlations between the consecutively decoded state pair frequencies *T* and the corresponding distances *D* were found even at replay speeds of about 14 items per second (i.e. inter-event intervals of 60-80ms, *r* = −0.018, permutation test: *r* = −0.003, t-test of sequence vs permutation results *t*_199_ = −4.42, *p* < .001, corrected for multiple comparisons; corresponding test for events separated by faster events at 40-60 ms: *p* = .18; *p* < .05 for all slower sequences; Fig. S5 and S6). This supports our conclusion that our findings in the resting-state data may reflect fast sequential replay in the human hippocampus.

In combination, these analyses show that sequences of hippocampal fMRI activity patterns during rest were systematically related to previous task experience. Interestingly, we found no such effect when we included backward distances between states instead of the forward distance in the model. This indicates that the sequences of hippocampal activity patterns became directionally structured in correspondence to participants’ task experience, and suggests that the hippocampus was engaged in forward replay in the post-task rest period.

Finally, we investigated the functional significance of hippocampal replay of abstract task states. One idea is that replay helps to form, or further solidify, a representation of the transitions between states of the task^23–25^. We therefore tested for a relationship between sequential state reactivation during rest and better representation of states during the task, as measured through cross-validated state decoding accuracy in fMRI data recorded during task performance. We did not find any evidence of a relationship between hippocampal sequenceness at rest and decoding of states during task performance (*r* = −.05, *p* > .05). However, we did find a significant correlation between hippocampal sequenceness at rest and state representations in the orbitofrontal cortex during the task (*r* = −.47, *p* = .005). Notably, in previous work we have shown that improved state decoding in the orbitofrontal cortex is associated with better decision making in this task, see^16^. Indeed, in the current dataset we also found a relationship between the change in orbitofrontal decoding accuracy during the task and improvements in task performance. That is, runwise decoding within the first session was correlated with runwise error rates (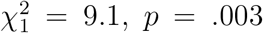, using the same decoding measure as used before, see ref. 16). This was not the case for on-task decoding in the hippocampus (*p* = .87, interaction with ROI: 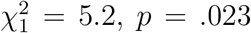). This finding therefore suggests a role for hippocampal replay in supporting the integrity of task-relevant orbitofrontal state representations during decision making. We also tested for a direct relationship between hippocampal replay at rest and behavioral measures of task performance, but did not find any evidence for a relationship between sequenceness and reaction times, error rates, or the change in these measures across runs (all *p*s > .10). Together, these results suggests that hippocampal replay supports the offline formation or maintenance of a ‘cognitive map’ of the task, while the orbitofrontal cortex is deployed to represent such a map during decision making^16,26^.

## Discussion

Our findings demonstrate that fMRI patterns recorded from the human hippocampus during rest reflect sequential replay of non-spatial task states previously experienced in a decision-making task. Previous studies have relied on sustained elevated fMRI activity in the hippocampus or sensory cortex as evidence for replay^18–20,27^, investigated wholebrain MEG signals^28^, or studied EEG sleep spindles and memory improvements that are thought to index replay activity^29–33^, but were not able to directly demonstrate sequential replay in the human hippocampus. Our study represents an important extension of these findings by providing evidence of sequential offline reactivation of non-spatial decision-making states in the human hippocampus.

Evidence of sequentiality and localization of replay in the human hippocampus is in direct correspondence with animal studies in which replay has been shown to be sequential and specific to hippocampal place cells, e.g.^34^. Importantly, unlike the majority of previous investigations in animals, the here reported sequences of activation patterns signify the replay of non-spatial, abstract task states. Our results therefore add to a growing literature proposing a significant role for ‘cognitive maps’ in the hippocampus in non-spatial decision making^3,8,26,35^.

Our findings are in line with the idea that the human hippocampus samples previous task experiences in order to improve the current decision-making policy, a mechanism that has been shown to have unique computational benefits for achieving fast and yet flexible decision making^23–25^. Dating back to Tolman^36^, this idea requires a neural mechanism that elaborates on and updates abstract state representations of the current task, regardless of the task modality. Several studies have suggested that the hippocampus and adjacent structures support a broad range of relational cognitive maps^35^, as evidenced by hippocampal encoding of not only spatial relations but also temporal^37,38^, social^7^, conceptual^6^ or general contingency relations^39^. Here, we showed that the human hippocampus not only represents these abstract task states, but also performs sequential offline replay of these states during rest.

Our results imply a relationship between hippocampal replay and the representation of decision-relevant task states that are thought to reside in the orbitofrontal cortex^16,22,40–42^. The relationship between ‘offline’ hippocampal sequenceness and the fidelity of ‘online’ orbitofrontal task-state representations raises the possibility that the hippocampus supports the maintenance and consolidation of state transitions that characterize the task and are employed during decision making^38^. Given our findings—and recent evidence implicating hippocampal place and entorhinal grid cells in signaling non-spatial task-relevant stimulus properties^6,8^—a crucial question for future studies will be to further specify how flexible, task specific representations in the hippocampus interact with task representations in other brain regions^26^. Of particular interest will be investigations asking whether neural populations in the hippocampus and entorhinal cortex share a common neural code for abstract task states with orbitofrontal^16^ and medial prefrontal regions^43^, as indicated by recent studies^6,44,45^. Together with our findings, this research promises to shed light on the neural representations and computations underlying memory and decision making.

## Experimental Procedures

### Participants

Thirty nine participants were selected according to standard fMRI screening criteria (right handedness, 18–35 years of age, normal or corrected-to-normal vision and no contraindication for fMRI) from the Princeton University community, and were compensated with $20 per hour plus up to $5 performance-related bonus. Six participants were excluded from analysis due to either technical errors (3 participants), violation of performance criteria standards (2 subjects with over 3 times the average error rate in the last two blocks of the experiment) or incomplete data (1 participant). The final sample consisted of 33 participants (22 female, mean age 23.4 years). All participants provided informed consent and the study was approved by Princeton University’s Institutional Review Board.

### Stimuli

Stimuli consisted of spatially superimposed images of a face and a house (see^16^; face images from http://faces.mpdl.mpg.de/faces described in^46^, see Fig. 1). Faces and houses could be classified as either young or old, e.g. a stimulus could show an old face image blended with a contemporary (i.e., young) house image. Four classes of stimuli were possible: (1) two old or (2) two young face and house pictures, (3) a young face with an old house or (4) vice versa.

### Task

The task was identical to Schuck et al. 2016 and will be described only briefly. Each trial began with the display of the mapping of a left and right key to a young and old response (changing randomly trialwise) below a fixation cross for 1.2s (range: 0.5–3.5s). Then, a compound face-house stimulus was shown for 3.3s (range: 2.75-5s; Fig. 1) and participants had to make an age judgment about one of the two image categories. Participants knew which category of the stimulus they had to judge by applying the following rules: 1. before the first trial of each run, the category to judge was displayed on the screen; 2. Once the age of the relevant category changed (e.g., from young to old), the judged category changed on the next trial. 3. No age comparison was necessary on the first trial after a category change, i.e. each category was judged for at least two trials in a row before a switch. The average trial duration was 4.5s (range: 3.25-8.5s), all timings were randomly drawn from a truncated exponential distribution and the response deadline was 2.75s. The category instruction cue at the beginning of a run was displayed for 4s. Erroneous or time-out responses led to feedback (written above stimulus for 0.7s) and trial repetition. If an error trial involved an age change (and thus would require a category switch on the next trial), participants had to repeat the trial before the error as well as the error trial, giving them the chance to observe the age change. Otherwise, they had to repeat the trial on which they made the error.

### Design

Participants underwent two fMRI sessions. The first session began with the display of written instructions while participants underwent a functional scan (group 1), or a 5 minute resting-state scan followed by instructions (group 2). The instructions explained the rules of the task and contained a training phase in which simple age judgments had to be made on (non-overlapping) face and house images. The images shown in this period were later used in the task, thus familiarizing participants with the age judgment aspect of the task as well as the stimuli. The instructions furthermore involved an annotated walk-through of four trials of the real task (i.e., with overlapping images and the requirement to switch attention after an age change). Following the instructions, participants performed 4 runs of the task (97 trials per run, 388 total). Each run lasted about 7-10 minutes and participants were given the chance to rest briefly between runs. A 5 minute fieldmap scan was done between runs 2 and 3, resulting in a longer break for participants. After run 4, participants underwent a resting state scan as well as a structural scan. Lights were turned off during resting-state scans and participants were instructed to stay awake for the entire duration of the scan (5 minutes, 100 TRs). The second session was identical for all participants and involved the following scans: resting state, 2 task runs, fieldmap, 2 task runs, resting state and structural scan. Thus, overall, participants performed 8 task runs and 3 (group 1) or 4 (group 2) resting-state scans. In all other regards, the task design involved the same characteristics as detailed in Schuck et al. (2016).

### Behavioral Analyses

Behavioral analyses were done using mixed effects models implemented in the package lme4^47^ in R^48^. The model included fixed effects for Block, Condition, Category and intercept. Participants were considered a random effect on the intercept and the slopes of all fixed effects. The reported *p*-values correspond to Wald chi-square (*χ*^2^) tests as implemented in R. Reaction time (RT) analyses were done on error-free trials only and reflect the median RT within each factor cell.

### fMRI Scanning Protocol

Magnetic-resonance images were acquired using a 3-Tesla Siemens Prisma MRI scanner (Siemens, Erlangen, Germany) located at the at the Princeton Neuroscience Institute. A T2*- weighted echo-planar imaging (EPI) pulse sequence was used for functional imaging (2×2 mm in plane resolution, TR = 3000 ms, TE = 27 ms, slice thickness = 2 mm, 53 slices, 96×96 matrix (FOV = 192 mm), iPAT factor: 3, flip angle = 80°, A→P phase encoding direction). Slice orientation was tilted 30° backwards relative to the anterior – posterior commissure axis to improve acquisition of data from the orbitofrontal cortex^49^. Field maps for distortion correction were acquired using the same resolution (TE1 = 3.99ms) and a MPRAGE pulse sequence was used to acquire T1-weighted images (voxel size = 0.9^3^ mm). The experiment began 20 seconds after acquisition of the first volume of each run to avoid partial saturation effects.

### fMRI Data Preprocessing

FMRI data preprocessing was done using SPM8 (http://www.fil.ion.ucl.ac.uk/spm) and involved fieldmap correction, realignment, and co-registration to the segmented structural images. The task data used to train the classifier were further submitted to a mass-univariate general linear model that involved run-wise regressors for each state (see below), nuisance regressors that reflected participant movement (6 regressors) and run-wise intercepts. The resulting voxelwise parameter estimates were z-scored and spatially smoothed (4 mm FWHM). The resulting activation maps were used as the training set for a support-vector. machine with a radial basis function (RBF) kernel that was trained to predict the task state from which a particular activation pattern came from LIBSVM^50^. Like the activation maps used for classifier training, the resting-state data were z-scored and smoothed (4mm FWHM). Anatomical ROIs were created using SPM’s `wfupick` toolbox. The hippocampus (HC) was defined as the left and right hippocampus AAL labels. The orbitofrontal cortex was defined as in^16^. Behavioral analyses and computations within the assumed graphical model of state space (see below) were done using R^48^.

### fMRI Classification Analysis

The support vector machines were trained on 8 maps of parameter estimates (“betas”) for each of the 16 states (one map for each state and run) restricted to the anatomical mask of the hippocampus (back-transformed into each subject’s individual brain space) or the orbitofrontal cortex. Classification accuracy was assessed with a leave-one-run-out cross-validation scheme in which data from 7 runs were used for training and the held-out run was used for testing (Fig. 2). The resting-state analysis used a classifier trained on all available task data (8 runs). This classifier was applied to each volume of the resting-state data and the most highly classified state was considered as the output of the classifier for that volume. The resulting sequence of predictions was the main focus of our analyses (see below). We obtained the distance to the hyperplane by dividing the decision value by the norm of the weight vector w, as specified in the libSVM webpage (http://www.csie.ntu.edu.tw/~cjlin/libsvm/faq.html#f4151). For each volume, we then calculated the average of the distances of all pairwise comparisons of the predicted class against all other classes, to obtain a proxy of how certain the classifier is in its prediction. Student t tests pertaining to decoding results were one-tailed, given the *a priori* expectation of larger-than-chance decoding in the hippocampus.

### Sequenceness Analysis

The main question of the sequenceness analysis was whether the state transitions decoded from resting state scans, *T*, were related to the distance between states experienced during the task, *D*. To this end, we analyzed the neural state transitions *T* with logistic mixed-effects models, using the lme4^47^ package in R^48^. Because the slow hemodynamic response function leads to encoding of sequential structure in activity patterns (i.e., there is high similarity between temporally adjacent patterns), a classifier trained on sequential task data can be biased to decode states in a similar sequence to the training data, even if the test data are random (i.e., the ‘sequenceness’ identified in the test data comes from the training data, not the test data). We therefore applied the trained classifier to matched fMRI noise (see below) and used the resulting spurious ‘state transitions,’ *T*(*ϵ*), as a covariate that would account of the spurious base rate of transitions that is due to the classifier rather than the data. Applying these models to control conditions consistently yielded non-significant effects of sequenceness, showing that this analysis appropriately controls for the above mentioned spurious structure that is observable for instance in the significant correlations between *D* and *T* in the noise data (Fig. 3F). Specifically, our model included the following fixed effects: (1) the distance between states, *D*, which was the regressor of interest, and as regressors of no interest (2) the transition probabilities obtained in the above mentioned noise simulations, *T*(*ϵ*), (3) an orthogonal quadratic polynomial of *T*(*ϵ*) that was included in order to account for as much noise-related variance as possible, and (4) an intercept. Models of change in sequenceness across PRE, INSTR and TASK conditions (Fig. 4) additionally involved interaction terms of condition with the distance *D* and condition with the noise transitions *T*(*ϵ*). Participant identity was included as a random factor to account for between subject variability. To capture state-related variability (state frequency effects affect the distribution of state transitions), state identity *s_j_* of a transition from state *i* to state *j* was used as an additional random effect nested within subject. Participant and state were random grouping factors for all fixed effects with exception of the quadratic expansion of *T*(*ϵ*), where including these random factors caused problems in fitting the logistic regression models.

Formally, the model followed the general assumption that the number of transitions *Y* is drawn from a binomial distribution of *n* draws and probability *T*:

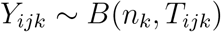

where *n_k_* corresponds to the number of measurements for subject *k*, and *i* and *j* index the outgoing and incoming states of a given transition. The logit transformed probabilities *T* (shown in Fig. 2D; logit is the canonical link function for binomial models) were then modeled in a mixed effects regression model with the above mentioned fixed and random effects structure:

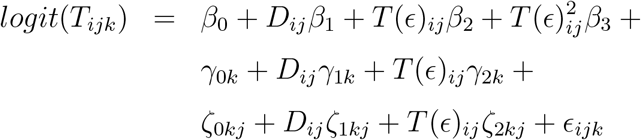

In the text, we describe the fixed effect of *D, β*_1_ in the models, as ‘sequenceness,’ and the fixed effect of *T*(*ϵ*), *β*_2_, as ‘randomness’ (Fig. 4B,C). The subject-specific random effects of *D, γ*_1*k*_, were used as individual sequenceness indicators in the correlations in Fig. 4F,G. The state- and subject-specific random effects are indicated by *ζ*. Correlations between random effects were estimated. Model comparisons were conducted using likelihood-ratio tests by comparing models that included the noise transitions *T*(*ϵ*) with versus without the fixed effect regressor of distance (sequenceness), or without the condition interaction terms to the full models that included these terms. The random effects structure was kept constant across these comparisons.

T-tests pertaining to sequenceness results (number of steps, etc.) are one-tailed, given our *a priori* expectation of larger sequenceness in the hippocampus compared to the various controls.

### Alternative Task Transition Matrices

As mentioned in the main text, for reasons of model comparability we used the 1-step transitions of the task as a basis to test alternative replay models. The 1-step transition matrix simply reflects from which state one could proceed to which other states in one trial. The alternative task transition matrices were based on the assumption that the hippocampus has access to only partial state information, and hence correspond to transition matrices defined over subsets of states.

For instance, to compute the transition matrix of the “stimulus model” we defined 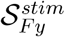 as the subset of states in which Fy was the stimulus:

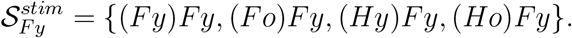

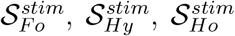 were the corresponding subsets of states in which Fo, Hy and Ho were the stimuli, respectively. The 1-step distance matrix was then computed such that every transition between two states *s_i_* and *s_j_* in the complete task state diagram was converted into four transitions from *s_i_* to all four states that are part of the same subset as *s_j_*, that is 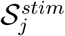. All resulting transitions are summed, and normalized so that all exiting transitions from a state would sum to 1. The new transition between 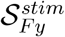 and 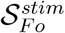 would therefore count within it (*Fy*)*Fy* → (*Fy*)*Fo*, (*Ho*)*Fy* → (*Fy*)*Fo* and (*Hy*)*Fy* → (*Fy*)*Fo*, whereas the new transition between 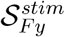 and 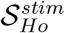 would only count within it (*Fo*)*Fy* → (*Fy*)*Ho*. After normalization, these would be 3/8 and 1/8, respectively.

For the other alternative models, we defined subsets of states that have the same current attended category, and subsets of states that have the same current and previous attended categories, and then computed the transition matrices as described above. The 1-step transition matrices of these alternative models are shown in Figure 5A-C. The reverse replay transition matrix was simply the transpose of the full task 1-step transition matrix.

### Synthetic fMRI Data and Noise Simulations

In order to estimate to what extend training the classifiers on sequential data influenced the sequenceness of their predictions, we simulated, for each participant, individually spatio-temporally matched fMRI noise, and applied the classifiers to these data. For each participant and resting state session, we first extracted fMRI data from the hippocampus and the orbitofrontal cortex. As in the classification analyses, we applied linear detrending to each voxel. We then estimated the average standard deviation of the voxels within these regions, as well as the average autocorrelation using an AR(1) model in R. Next, we used the neuRosim toolbox in R^51^ to simulate fMRI noise with the same standard deviation and temporal autocorrelation as the real data. Finally, we used AFNI’s 3dFWHMx and 3dBlurToFWHM functions to first estimate the spatial smoothness of the real data, and then smooth the simulated noise until it has the same effective smoothness. For each existing resting-state run, matched noise data with the same number of TRs and voxels were generated. Figure S1 (SI) shows the temporal and spatial smoothness of the real and simulated data separately for each run. In all cases, the properties of the simulated data did not differ from the real data, paired t-tests, all *p*s > .05.

Finally, we applied each participant’s classifier to the matched noise data. The classifier was identical to the classifier that was used in estimating the sequences of states from the real data. The resulting sequence of predicted states reflects the bias of the classifier to make sequential predictions because of pattern overlap in the training set, even when applied to noise, as well as any tendency of the classifier to decode certain states more often than others. We therefore used the sequence of states from this analysis to construct the nuisance covariate for the mixed effects models, i.e. the noise ‘transition matrix,’ *T*(*ϵ*), and to perform the appropriate comparisons in the correlation analysis. These comparisons between sequenceness in real data versus simulated noise in the correlation and mixed effect analyses indicated that the noise sequenceness *T*(*ϵ*) indeed explained a significant amount of sequential variability of the decoded states (see Figs. 3F,G, 4B, D), and thus served as a powerful control. Together with the permutation tests (Fig. 3B-D, 3F,G), the comparisons across brain regions (Fig. 4E) and the within-participant comparisons between the PRE, INSTR and TASK conditions (3B-D, H and 4A-D), our approach represents a stringent control of potential biases.

## Data Avalability

All raw fMRI data, and the sequence of decoded states used to generate results in Figures 2–5 will be made publicly available upon publication.

## Code Avalability

All code used to generate results in Figures 2–5 will be made publicly available upon publication.

## Author Contributions

NWS, and YN designed research. NWS conducted research. NWS and YN analyzed and interpreted the data and wrote the manuscript.

## Acknowledgments

This research was funded by NIH grant R01DA042065 and funding from John Templeton Foundation. The opinions expressed in this publication are those of the authors and do not necessarily reflect the views of the John Templeton Foundation or those of NIH. We thank Nathaniel Daw, Christian Doeller, Eran Eldar and David Tank for helpful comments on this research and/or previous versions of the manuscript.

## References

1. Scoville, W. & Milner, B. Loss of recent memory after bilateral hippocampal lesions. Journal of Neurology, Neurosurgery & Psychiatry 20, 11–21 (1957).

2. Squire, L. R. Memory and the Hippocampus: A Synthesis From Findings With Rats, Monkeys, and Humans. Psychological Review 99, 195–231 (1992).

3. Eichenbaum, H, Dudchenko, P, Wood, E, Shapiro, M & Tanila, H. The hippocampus, memory, and place cells: is it spatial memory or a memory space? Neuron 23, 209–26 (1999).

4. O’Keefe, J. & Nadel, L. The hippocampus as a cognitive map (Oxford University Press, Oxford, 1978).

5. McKenzie, S. et al. Hippocampal representation of related and opposing memories develop within distinct, hierarchically organized neural schemas. Neuron 83, 202–215 (2014).

6. Constantinescu, A. O., OReilly, J. X. & Behrens, T. E. J. Organizing conceptual knowledge in humans with a gridlike code. Science 352, 1464–1468 (2016).

7. Tavares, R. M. et al. A Map for Social Navigation in the Human Brain. Neuron 87, 231–243 (2015).

8. Aronov, D., Nevers, R. & Tank, D. W. Mapping of a non-spatial dimension by the hippocampal-entorhinal circuit. Nature 543, 719–722 (2017).

9. Foster, D. J. & Knierim, J. J. Sequence learning and the role of the hippocampus in rodent navigation. Current Opinion in Neurobiology 22, 294–300 (2012).

10. Wikenheiser, A. M. & Redish, A. D. Decoding the cognitive map: Ensemble hippocampal sequences and decision making. Current Opinion in Neurobiology 32, 8–15 (2014).

11. Skaggs, W. E. & McNaughton, B. L. Replay of neuronal firing sequences in rat hippocampus during sleep following spatial experience. Science 271, 1870–1873 (1996).

12. Johnson, A. & Redish, D. Neural ensembles in CA3 transiently encode paths forward of the animal at a decision point. Journal of Neuroscience 27, 12176–89 (2007).

13. Karlsson, M. P. & Frank, L. M. Awake replay of remote experiences in the hippocampus. Nature Neuroscience 12, 913–918 (2009).

14. Carr, M. F., Jadhav, S. P. & Frank, L. M. Hippocampal replay in the awake state: a potential substrate for memory consolidation and retrieval. Nature Neuroscience 14, 147–53 (2011).

15. Girardeau, G., Benchenane, K., Wiener, S. I., Buzsáki, G. & Zugaro, M. B. Selective suppression of hippocampal ripples impairs spatial memory. Nature Neuroscience 12, 1222–1223 (2009).

16. Schuck, N. W., Cai, M. B., Wilson, R. C. & Niv, Y. Human Orbitofrontal Cortex Represents a Cognitive Map of State Space. Neuron 91, 1402–1412 (2016).

17. Tong, S. & Chang, E. Support vector machine active learning for image retrieval in Proceedings of the ninth ACM international conference on Multimedia October (ACM Press, New York, New York, USA, 2001), 107–118. doi:10.1145/500156.500159.

18. Peigneux, P. et al. Are Spatial Memories Strengthened in the Human Hippocampus during Slow Wave Sleep? Neuron 44, 535–545 (2004).

19. Deuker, L. et al. Memory Consolidation by Replay of Stimulus-Specific Neural Activity. Journal of Neuroscience 33, 19373–19383 (2013).

20. Staresina, B. P., Alink, A., Kriegeskorte, N. & Henson, R. N. Awake reactivation predicts memory in humans. Proceedings of the National Academy of Sciences of the United States of America 110, 21159–64 (2013).

21. Axmacher, N., Elger, C. E. & Fell, J. Ripples in the medial temporal lobe are relevant for human memory consolidation. Brain 131, 1806–1817 (2008).

22. Wilson, R. C., Takahashi, Y. K., Schoenbaum, G. & Niv, Y. Orbitofrontal cortex as a cognitive map of task space. Neuron 81, 267–79 (2014).

23. Sutton, R. S. Integrated architectures for learning, planning, and reacting based on approx dynamic prog in Proceedings of the 7th International Conference on Machine Learning (1990), 216–224. doi:10.1.1.51.7362.

24. Van Seijen, H. & Sutton, R. S. A Deeper Look at Planning as Learning from Replay. Proceedings of the 32nd International Conference on Machine Learning 37 (2015).

25. Russek, E. M., Momennejad, I., Botvinick, M. M. & Gershman, S. J. Predictive representations can link model-based reinforcement learning to model-free mechanisms. bioRxiv 083857. doi:10.1101/083857 (2016).

26. Kaplan, R., Schuck, N. W. & Doeller, C. F. The Role of Mental Maps in Decision-Making. Trends in Neurosciences 40, 256–259 (2017).

27. Bergmann, T. O., Mölle, M., Diedrichs, J., Born, J. & Siebner, H. R. Sleep spindle-related reactivation of category-specific cortical regions after learning face-scene associations. NeuroImage 59, 2733–2742 (2012).

28. Kurth-Nelson, Z., Economides, M., Dolan, R. J. & Dayan, P. Fast sequences of non-spatial state representations in humans. Neuron 91, 194–204 (2016).

29. Cox, R., Hofman, W. F., de Boer, M. & Talamini, L. M. Local sleep spindle modulations in relation to specific memory cues. NeuroImage 99, 103–110 (2014).

30. Rasch, B., Buchel, C., Gais, S. & Born, J. Odor Cues During Slow-Wave Sleep Prompt Declarative Memory Consolidation. Science 315, 1426–1429 (2007).

31. Antony, J. W., Gobel, E. W., O’Hare, J. K., Reber, P. J. & Paller, K. A. Cued memory reactivation during sleep influences skill learning. Nature Neuroscience 15, 1114–1116 (2012).

32. Siapas, A. G. & Wilson, M. A. Coordinated interactions between hippocampal ripples and cortical spindles during slow-wave sleep. Neuron 21, 1123–1128 (1998).

33. Ji, D. & Wilson, M. A. Coordinated memory replay in the visual cortex and hippocampus during sleep. Nature Neuroscience 10, 100–107 (2007).

34. Lee, A. K. & Wilson, M. A. Memory of Sequential Experience in the Hippocampus during Slow Wave Sleep. Neuron 36, 1183–1194 (2002).

35. Schiller, D. et al. Memory and Space: Towards an Understanding of the Cognitive Map. Journal of Neuroscience 35, 13904–13911 (2015).

36. Tolman, E. C. Cognitive maps in rats and men. Psychological Review 55, 189–208 (1948).

37. Fortin, N. J., Agster, K. L. & Eichenbaum, H. B. Critical role of the hippocampus in memory for sequences of events. Nature Neuroscience 5, 458–62 (2002).

38. Schapiro, A. C., Kustner, L. V. & Turk-Browne, N. B. Shaping of object representations in the human medial temporal lobe based on temporal regularities. Current Biology 22, 1622–1627 (2012).

39. Stachenfeld, K. L., Botvinick, M. M. & Gershman, S. J. The hippocampus as a predictive map. Nature Neuroscience 20, 1643–1653 (2017).

40. Bradfield, L. A., Dezfouli, A., van Holstein, M., Chieng, B. & Balleine, B. W. Medial Orbitofrontal Cortex Mediates Outcome Retrieval in Partially Observable Task Situations. Neuron, 1–13 (2015).

41. Howard, J. D. & Kahnt, T. Identity-specific reward representations in orbitofrontal cortex are modulated by selective devaluation. Journal of Neuroscience 37, 3473–16 (2017).

42. Nogueira, R. et al. Lateral orbitofrontal cortex anticipates choices and integrates prior with current information. Nature Communications 8, 14823 (2017).

43. Schuck, N. W. et al. Medial Prefrontal Cortex Predicts Internally Driven Strategy Shifts. Neuron 86, 331–340 (2015).

44. Doeller, C. F., Barry, C. & Burgess, N. Evidence for grid cells in a human memory network. Nature 463, 657–61 (2010).

45. Jacobs, J et al. Direct recordings of grid-like neuronal activity in human spatial navigation. Nature Neuroscience 16, 1188–1190 (2013).

46. Ebner, N. C., Riediger, M. & Lindenberger, U. FACES - A database of facial expressions in young, middle-aged, and older women and men: Development and validation. Behavior Research Methods 42, 351–362 (2010).

47. Bates, D., Mächler, M., Bolker, B. & Walker, S. Fitting Linear Mixed-Effects Models Using lme4. Journal of Statistical Software 67, 51 (2015).

48. R Core Team. R: A Language and Environment for Statistical Computing Vienna, Austria, 2015.

49. Deichmann, R, Gottfried, J., Hutton, C & Turner, R. Optimized EPI for fMRI studies of the orbitofrontal cortex. NeuroImage 19, 430–441 (2003).

50. Chang, C.-c. & Lin, C.-j. LIBSVM 2011. doi:10.1145/1961189.1961199.

51. Welvaert, M., Durnez, J., Moerkerke, B., Verdoolaege, G. & Rosseel, Y. neuRosim: An R Package for Generating fMRI Data. Journal of Statistical Software 44, 1–18 (2011).

